# Cross-organ transcriptomic comparison reveals universal factors during maturation

**DOI:** 10.1101/2021.08.03.454962

**Authors:** Sandeep Kambhampati, Sean A. Murphy, Hideki Uosaki, Chulan Kwon

## Abstract

Various cell types can be derived from stem cells. However, these cells are immature and do not match their adult counterparts in functional capabilities, limiting their use in disease modeling and cell therapies. Thus, it is crucial to understand the mechanisms of maturation in vivo. However, it is unknown if there are genes and pathways conserved across organs during maturation. To address this, we performed a time-series analysis of the transcriptome of the mouse heart, brain, liver, and kidney and analyzed their trajectories over time. In addition, gene regulatory networks were reconstructed to determine overlapping expression patterns. Based on these, we identified commonly upregulated and downregulated pathways across all four organs. Key upstream regulators were also predicted based on the temporal expression of downstream genes. These findings suggest the presence of universal regulators during organ maturation, which may help us develop a general strategy to mature stem cell-derived cells in vitro.

## Introduction

Early mouse embryogenesis begins with fertilization and implantation, leading to the formation of a hollow egg cylinder by E5.0. During gastrulation, starting at E6.0, the three germ layers, endoderm, ectoderm, and mesoderm, are established (Takoaka et al. 2012, Tam et al. 1997, Robb et al. 2004). Gastrulation is followed by organogenesis at E8.0, during which these three germ layers rapidly proliferate and differentiate into specialized organ systems (Takoaka et al. 2012, Tam et al. 1997, Robb et al. 2004). Individual organs undergo organ-specific morphological changes, yet organogenesis is a coordinated process linked to the overall growth rate of the embryo.

This coordination can be achieved by genes with similar expression profiles in multiple organs. Genes can be broadly characterized into two groups based on their spatiotemporal expression patterns across multiple tissues. Housekeeping genes, which are expressed ubiquitously, serve to maintain basal cellular functions of a cell and are thus essential to general cell survival (Warrington et al. 2000). Selective genes, on the other hand, have enriched expression in several tissues (Liang et al. 2006). Transcriptional analysis has shown that these selective genes could mediate functional connectivity and organ-crosstalk among multiple organs (Qin et al. 2016). In the context of development, such crosstalk would be crucial to achieving coordinated organogenesis.

In particular, transcriptional dynamics are crucial for coordinating embryonic development. One shift in transcription occurs during embryonic development and is characterized by increased expression of genes with early organ-specific functions and decreased expression of genes controlling cell division and general morphogenesis (Cardoso-Moreira et al. 2019). In mice, which are precocial, the second shift overlaps with birth and is characterized by increased expression of genes with late organ-specific function and again, decreased expression of genes controlling cell division and general morphogenesis (Cardoso-Moreira et al. 2019).

It may therefore be expected that transcriptional downregulation may be similar and coordinated across multiple organs, especially with regards to the decrease in proliferation that is generally characteristic of organogenesis and maturation. Furthermore, it can be hypothesized that transcriptional up-regulation may follow certain similarities across multiple organs as well. From an evolutionary perspective, new regulatory mechanisms would develop by building upon existing ones. So as new organs evolved, their mechanisms of growth and development would likely share many similarities (Goss 1990). Indeed, there are many common aspects of growth among different organs, such as the impact of use and disuse on promoting positive vs negative growth, hypertrophy and hyperplasia, and metabolic rate (Goss 1990). Hormones that circulate through the bloodstream could provide a mechanism to coordinate changes in metabolism and growth of multiple organs throughout development.

We thus hypothesized that there may be common regulators of multiple organ development. To test this hypothesis, we analyzed the transcriptomes of hearts, brains, livers, and kidneys profiled at various developmental stages. The heart and kidney originate from the mesoderm, the liver originates from the endoderm, and the brain originates from the ectoderm. The four organs chosen thus represent all three germ layers. While the development of these four organs differs due to organ-specific functions, there are several important similarities. The development of these organs at early stages between E8 to E13 is characterized by proliferation and migration of cells as progenitor cells within these organs begin to form the overall structure of the organs (Takoaka et al. 2012). Postnatally, cells in these organs all undergo structural, functional, and metabolic maturation. These processes are directed towards obtaining important functional characteristics of adult cells. In the case of cardiomyocytes and neurons, electrophysiological maturation is crucial for organ function in adults, and functional coupling of these two cell types promotes mutual maturation (He et al. 2018, Lopaschuk et al. 2010, Oh et al. 2016). Moreover, major changes in metabolism and energy handling, with increases in mitochondrial biogenesis and mitochondrial complexity occur during the development and maturation of cardiomyocytes, hepatocytes, neurons, and podocytes (Agostini et al. 2016, Yuan et al. 2020, Lopaschuk et al. 2010, Nishikawa et al. 2014). While stem/progenitor or immature cells rely mainly on glycolysis for ATP production, which is necessary to meet their proliferation needs (Shyh-Chang et al. 2017), all four cell types switch to a primary reliance on oxidative phosphorylation for their energy needs by their adult, mature stages.

## Results

### Global transcriptomic maturation

A total of 479 brain, 147 liver, 142 kidney, and 212 heart murine microarray expression profiles, obtained from Gene Expression Omnibus, were combined to create a full organ dataset. The data for each organ represents wild-type and un-treated samples, covering the developmental stages of organogenesis and maturation. Data collected from timepoints between E8-E11 were classified as ‘Early’, E12-E15 as ‘Mid’, E16-birth as ‘Late’, P3-P10 as ‘Neonate’ and 2-3 months to 30 weeks as ‘Adult’.

We first applied Principal Component Analysis (PCA) to the combined dataset of all four organs. PCA is a dimensionality reduction algorithm that can be used to identify principal components, or eigenvectors of the data covariance matrix, that capture maximal variance of the data features. We see that by looking at the first 3 principal components (PCs) together, all 4 organs cluster together early in development, after which each follows their own unidirectional, non-branching trajectory (Figure 1a). PCA thus reveals that the differences between organs are more similar at early embryonic stages and diverge throughout the course of development and maturation. This aligns with the current view of organogenesis during development. Furthermore, plotting the principal components, PC2 vs PC3, we see that PC2 captures a common developmental trajectory between all 4 organs studied (Figure 1b). This indicates that there are common processes underlying the development and maturation of these organs, and therefore, potentially common regulators of maturation.

**Fig. 1.**
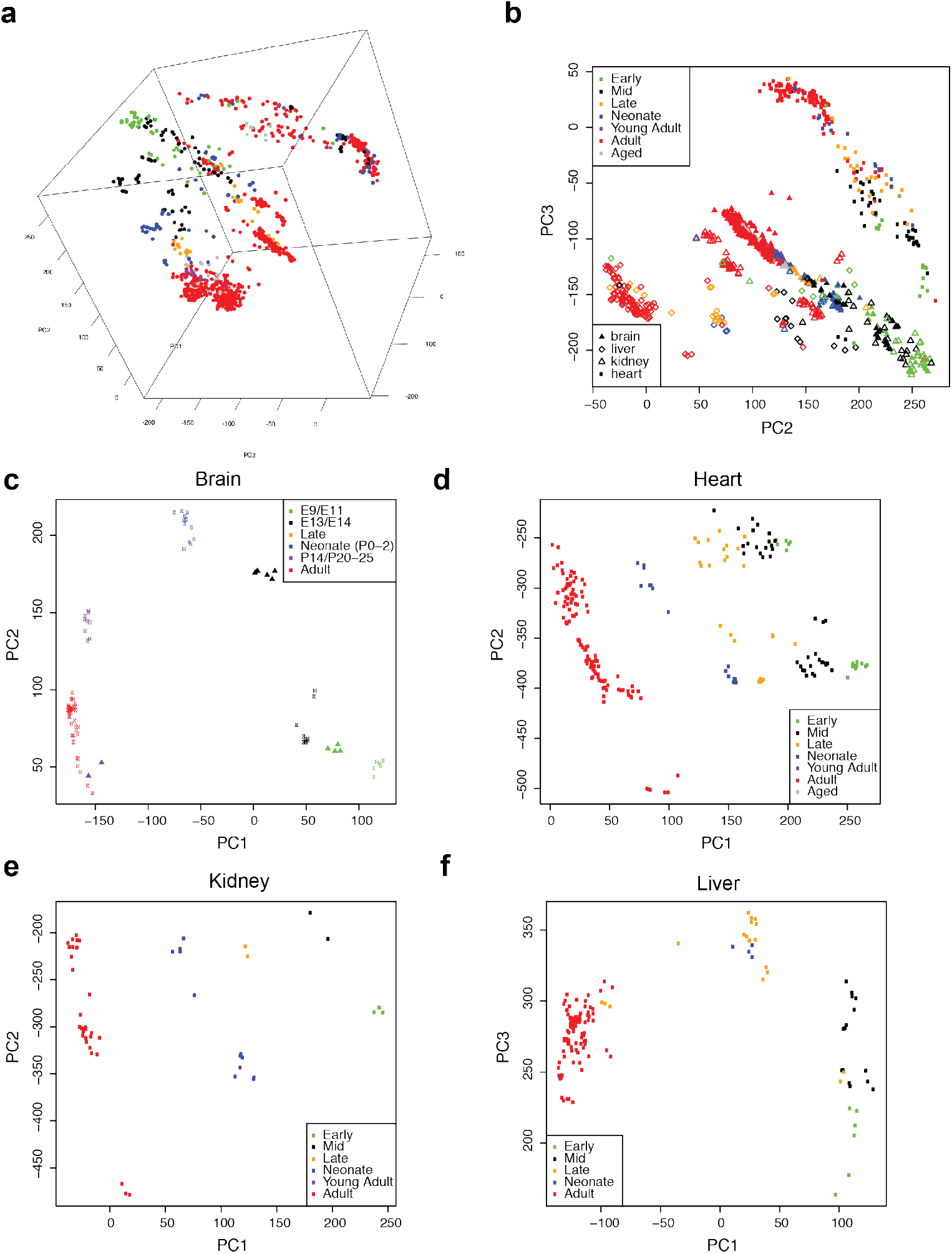
Cross-organ PCA clustering of transcriptomes. **a)** 3D PCA plot demonstrating 3 PCs simultaneously capturing development and maturation of the heart, liver, kidney, and brain. Color follows the same legend as (b). Heart: 213 microarray datasets. Early (E8-11, N=17, green), mid (E12-14, N=39, black), late (E16-18, N=26, orange) embryonic, postnatal (P3-10, N=16, blue) and adult (N=114, red). Brain: 479 datasets. Early (N=18), mid (N=38), Late (N = 12), neonate (N=32), adult (N=359). Kidney: 142 datasets Early (N=39), Mid (N=22), Late (N=2), Neonate (N=12), adult (N=67). Liver: 147 datasets. Early (N=6), Mid (N = 14), Late (N = 18), Neonate (N=4), adult (N = 104). **b)** PC2 vs PC3 of development and maturation of all 4 organs analyzed in this study. Principal component analysis revealed that differences between organs were minimal at the early embryonic stage and were more obvious at the later stages. As maturation seems to happen via a similar trajectory in all organs, we hypothesized that there could be common maturation processes and presumably common regulators of maturation through organs. **c,d,e,f)** Principal Component Analysis (PCA): PCA plots of single organ development. Single dot represents one microarray dataset. Samples were clustered and aligned through PC1 axis as maturation progress. Each individual organ shows a clear unidirectional trajectory from early stage to adult stage.

PCA of the data from each individual organ captures organ-specific developmental trajectories (Figure 1c-f), with data collected from various time-points clustered by developmental stage (early embryonic, mid embryonic, late embryonic, neonatal and adult). In the kidney and heart, the developmental trajectory is primarily captured by PC1. In the liver and brain, the developmental trajectory is captured by a combination of PC1 and PC2, with the difference in gene expression between early embryonic and adult stages driving the variance explained by PC1. Altogether, these results indicate that there is a common trajectory from immature to mature among the four organs as well as organ specific trajectories that add additional dimensions unique to each specialized tissue.

### Expression trends identify conserved pathways

After identifying similar trajectories underlying the development of all four organs, we sought to identify the genes driving these similarities. We reasoned that genes following similar gene expression time-course patterns between various organs may link functional connectivity between the organs. Such similarities could reflect responses to signaling molecules coordinating organogenesis and maturation. We performed fuzzy clustering of organ-specific gene expression time series data in order to identify genes with similar trajectories between organs using the MFuzz R package. For simplicity, we focused on two trajectories: up-regulated, where expression progressively increased at each time point, or down-regulated where expression progressively decreased at each time point. After selecting the strongest genes in the cluster based on membership score, we analyzed overlap between the up-regulated and down-regulated genes from different organs. We ran Gene Ontology analysis to look for functional enrichment of genes that were up- or down-regulated in multiple organs (Supplementary Table 1).

The genes that were down-regulated (Figure 2a-d) were largely involved in cell cycle processes, enriched for GO terms such as DNA Replication, Mitotic Cell Cycle, and Nuclear Chromosome Segregation (Figure 2e). This follows the understanding that as development and maturation progresses, cell division slows along with an increased focus on functional maturation. The genes that were up-regulated (Figure 3a-d) were largely involved in metabolism, enriched for GO terms such as Oxidative Phosphorylation, Mitochondrion, Monocarboxylic Acid Catabolic Process (Figure 3e). This follows the understanding that as the physiological demand on cells increases throughout development, they must alter their energy handling apparatuses.

**Fig. 2.**
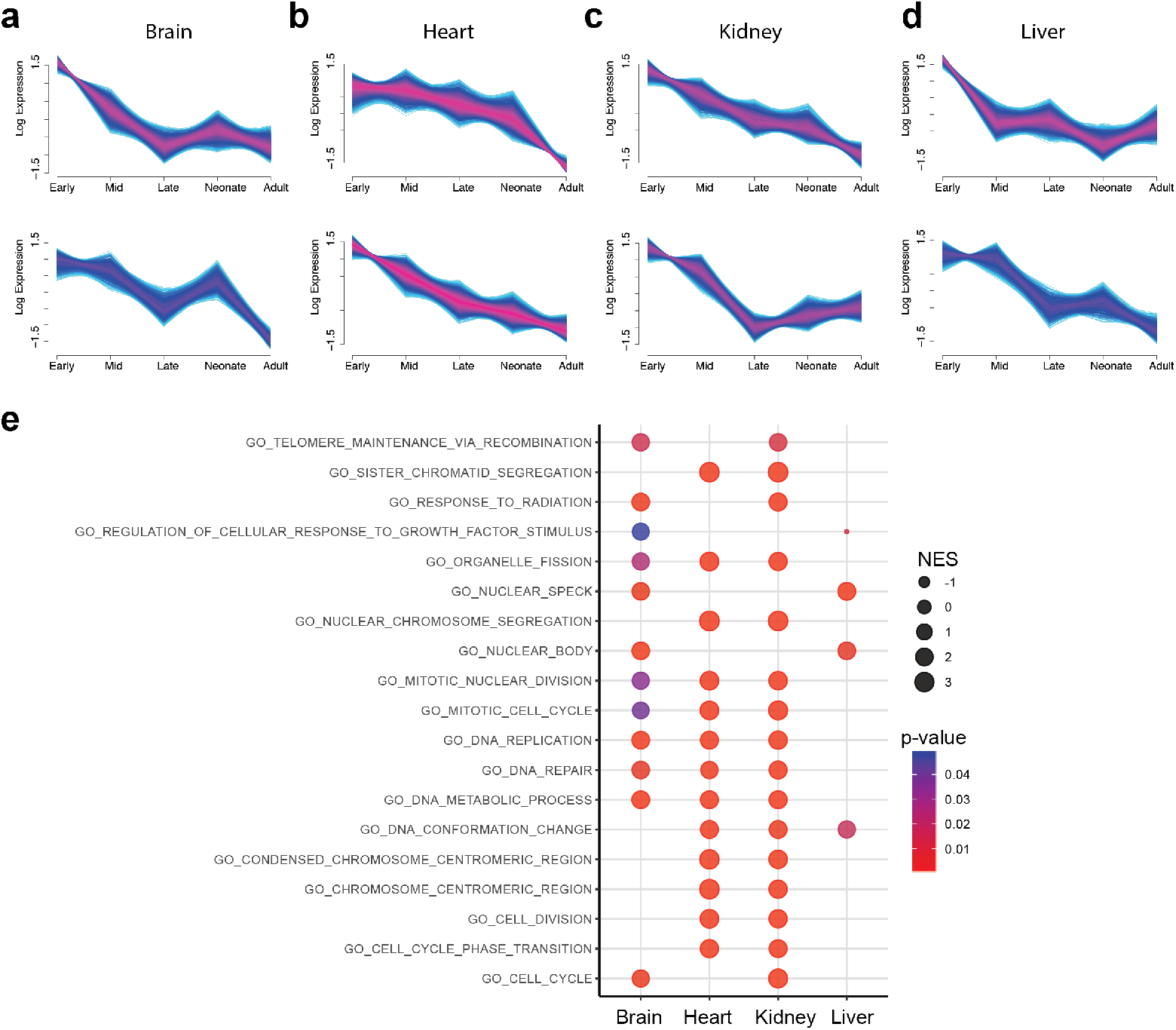
Fuzzy clustering of down-regulated genes over maturation. **a-d)** Fuzzy Clustering plots of gene expression trends using mfuzz. With this form of soft clustering, genes are assigned a membership score between 0 and 1 for each cluster, in contrast to hard assignment to a single cluster. Clusters for down-regulated genes are shown. Genes with cluster membership scores greater than 0.5 are shown, with lighter color indicating higher degree of cluster membership. **e)** Significant enriched GO terms for each organ shown, determined using genes with high membership scores (> 0.5) to down-regulated clusters. Color indicates p-value and size indicates normalized enrichment score. Those with overlap across organs include terms relevant to cell division.

**Fig. 3.**
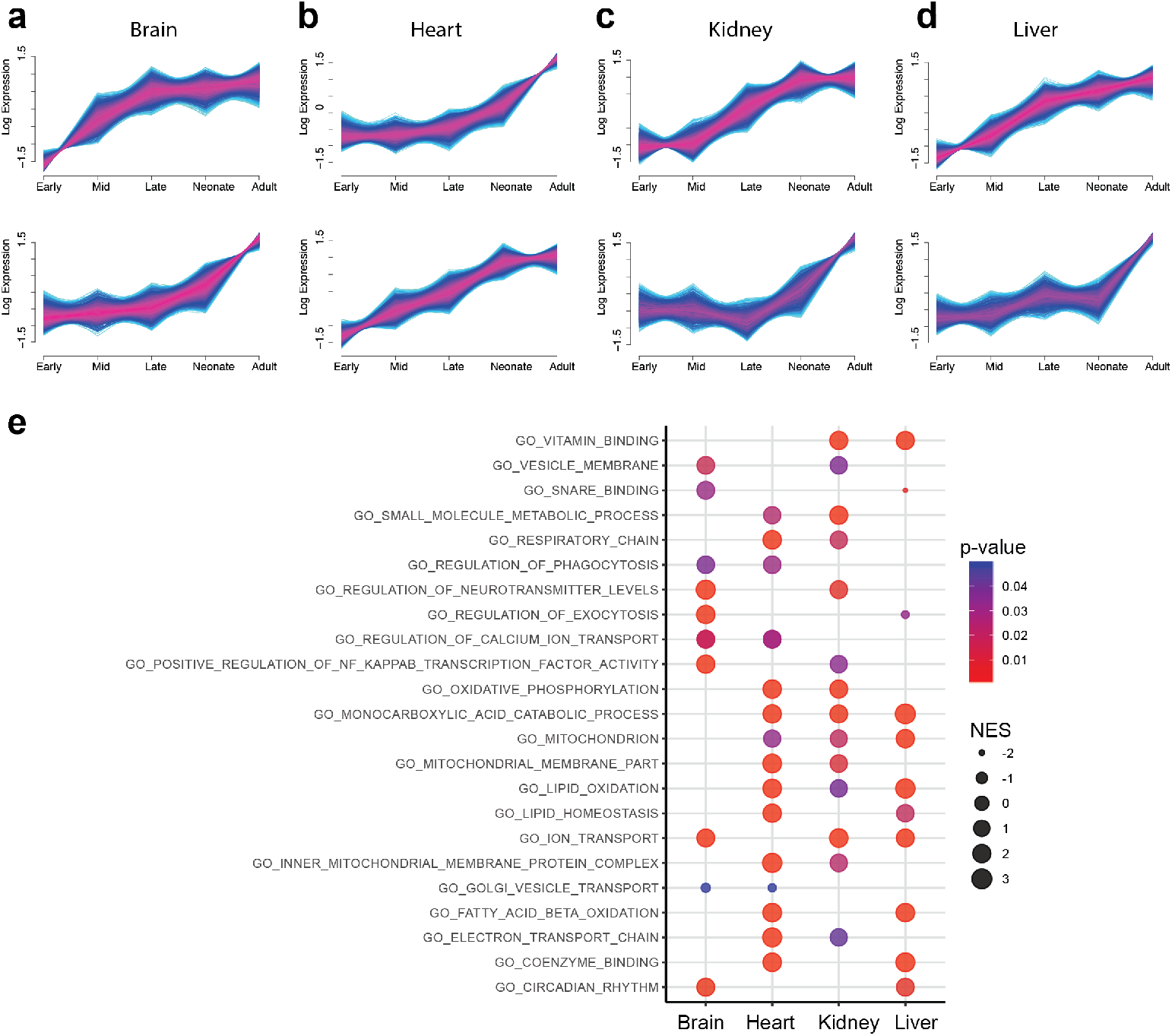
Fuzzy clustering of up-regulated genes over maturation. **a-d)** Genes with cluster membership scores greater than 0.5 are shown, with lighter color indicating higher degree of cluster membership. **e)** Significant enriched GO terms for each organ shown, determined using genes with high membership scores (> 0.5) to up-regulated clusters. Color indicates p-value and size indicates normalized enrichment score. Those with overlap across organs include terms relevant to metabolic processes.

### Gene regulatory network of multiorgan maturation

Given the similarities in up- and down-regulated genes during development across multiple organs, we next analyzed whether there were any similarities in functional organization of gene regulatory networks during development and maturation. Weighted gene coexpression network analysis was computed using the WGCNA R package. This identified modules of coexpressed genes in each organ (Figure 4c, 4f, 4i, 4l). We determined which modules were positively (Figure 4b, 4e, 4h, 4k) and negatively (Figure 4a, 4d, 4g, 4j) correlated with time by plotting the eigengene value at each timepoint. The eigengene is the first principal component of the gene expression matrix for a given module, and thus can be thought of as a weighted average representation of the gene expression profile for that module.

**Fig. 4.**
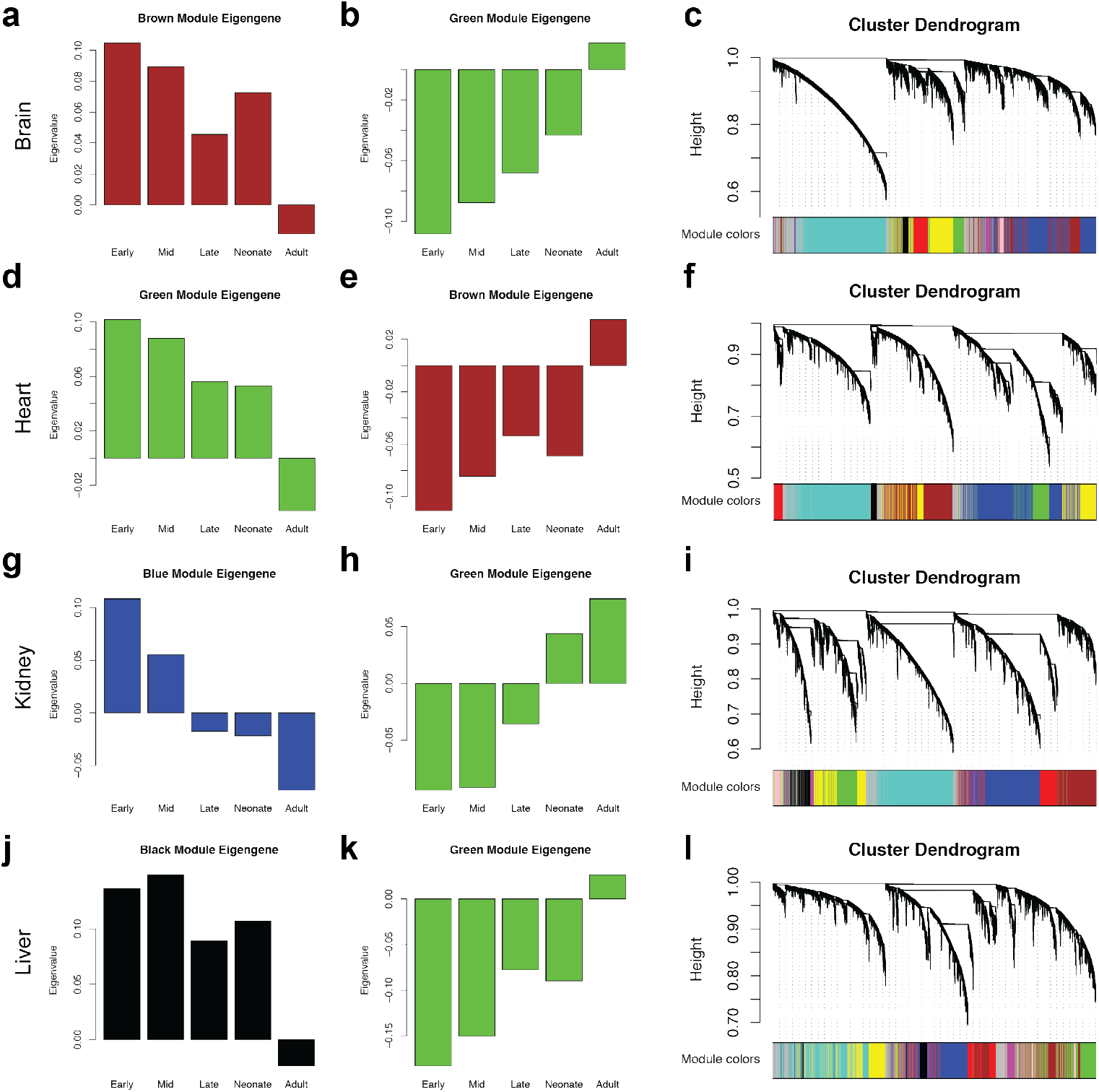
Weighted gene correlation network analysis of cross-organ maturation. **a-l)** On left, for each organ, the eigengene, a weighted average representation of the gene expression profile for that module, is shown for down- and up-regulated genes through development. The eigenvalue for a given eigengene is plotted at each developmental stage. On right, cluster dendrogram depicting reconstructed gene regulatory network for that organ. Branches within this dendogram correspond to different modules. Intramodular hub genes of these modules are located at the tips of the branches. WGCNA reveals strong functional organization within the gene regulatory networks controlling development and maturation. Enriched GO terms determined from genes strongly associated with up- and down-regulated modules are included in supplementary tables. In agreement with the results of fuzzy clustering, common terms associated with the up-regulated module across all four organs include metabolic terms while those with the down-regulated module include cell cycle terms.

The genes were compared among modules that increased in correlation with time. We found that Cdkn2, a cell cycle repressor, was found in all the positively correlated groups for all four organs (Supplementary Table 2). GO term analysis of each module showed many cell cycle terms in the negatively correlated modules and metabolic terms in the positively correlated modules (Supplementary Table 2). This indicates that metabolic and cell cycle suppression pathways become predominantly expressed in the adult cells. This was expected as cells slow their proliferation to maintain organ volume (Tumaneng et al. 2012).

### Prediction of upstream regulators of maturation

Next, based on the similarities in gene regulatory networks governing de velopment and maturation across the four organs, we sought to determine the common transcriptional regulators of maturation that may be driving these networks. We used Ingenuity pathway analysis (IPA) to generate z-scores for each upstream regulator through pairwise comparisons of gene changes between two adjacent timepoints. Differentially expressed genes between consecutive developmental stages were inputted to IPA giving us significantly enriched upstream regulators at each developmental timepoint. Positively enriched (activated) and negatively enriched (inhibited) upstream regulators were plotted across all timepoints and organs. This let us find transcription factors which are consistently regulated in maturation in multiple organs (Supplementary Table 3).

Positively enriched upstream regulators included cell cycle genes and regulators of proliferation and growth such as CDKN2A, CEBPA, and HDAC1/2 as well as regulators of metabolism such as the PPAR family (Figure 5a) These results are in accordance with the analysis using fuzzy clustering and gene regulatory network analysis and follow our understanding of rapid proliferation and metabolic changes as two critical processes of development and maturation. We would expect proliferation to be important earlier in development and metabolic changes to be important later in development; however, interestingly, activation of regulators from both of these categories tended to occur at similar stages of development.

**Fig. 5.**
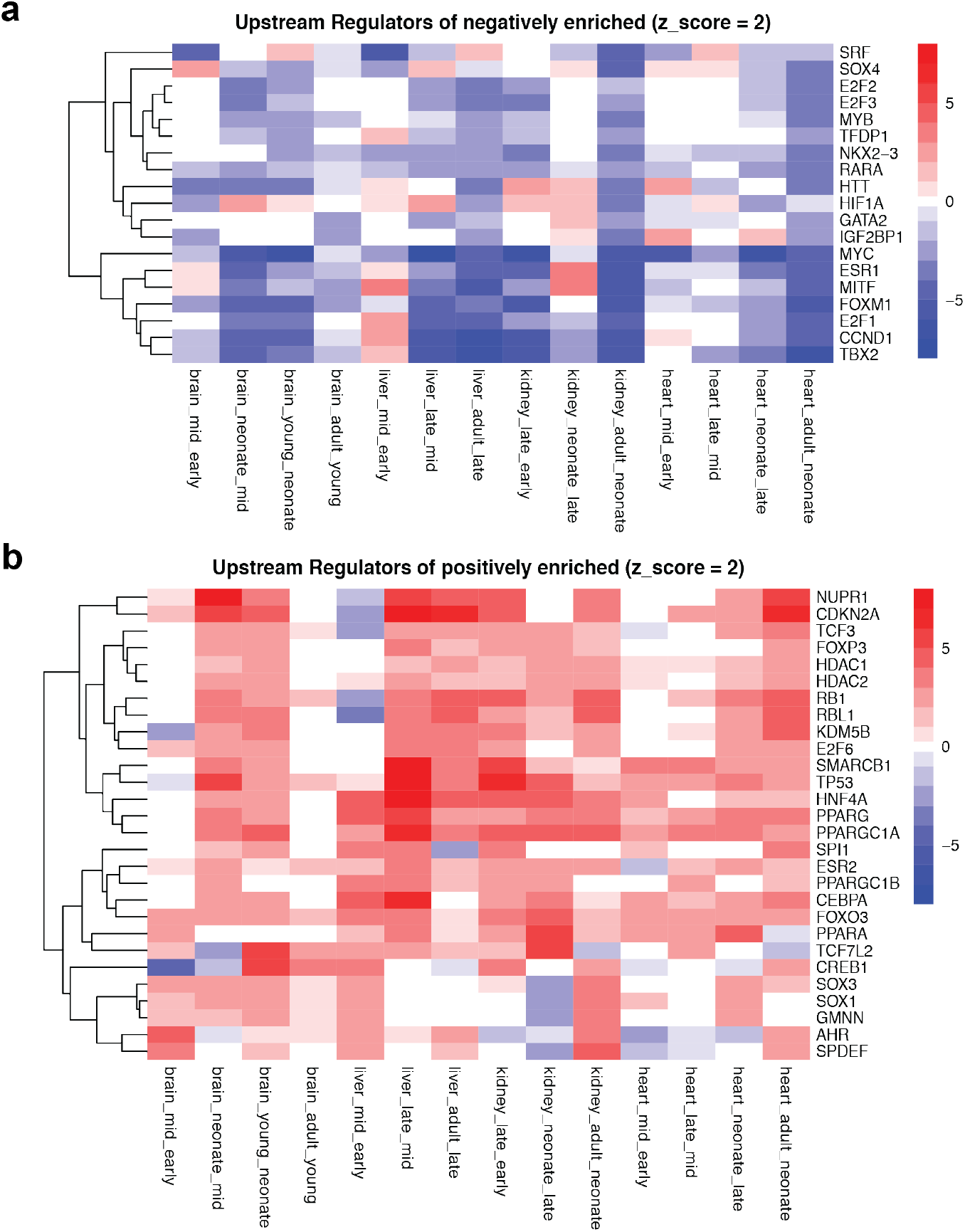
Predicted upstream regulators of global maturation. **(a-b)** Heatmap of the activation Z-scores of upstream transcriptional regulators, correspond to activation changes from one stage to the next. Red: Higher in later stage, Blue: Lower in later stage. Upstream regulators were identified by running IPA on differentially expressed genes between consecutive stages of development. More than 400 upstream regulators were involved in the maturation process of the organs analyzed. Among the regulators, 28 and 19 regulators were identified as commonly activated and inactivated ones.

Negatively enriched upstream regulators included genes involved with cell fate and development, such as GATA2, SOX4, and SRF among others (Figure 5b). Regulators involved in early fate decisions would naturally be inhibited as maturation progresses. Interestingly, several of these genes are strongly associated with development and maturation of single organs. For example, TBX2 is required for mesoderm differentiation. However, it showed strong negative enrichment in the brain (ectoderm) and kidney (endoderm) as well. Other genes, such as ESR1, IGF2BP1, and MYC are generally critical for processes such as cell proliferation and differentiation.

Altogether, the similarities in activated and inhibited transcriptional regulators include genes that might be expected due to their ubiquitous role in regulating gene expression and homeostatic processes. In addition, several genes that are well known for organ-specific roles appear to be important in other organs and germ layers, indicating a greater degree of functional connectivity among organs than previously appreciated.

## Discussion

In this study, we performed a meta-analysis of mouse microarrays to track gene expression during multiorgan maturation. A global trajectory of transcriptomic maturation was generated from PCA. We sought to capture the similarities in organ maturation through differential expression and gene regulatory network reconstruction. Although no single pathway appears to be a master regulator of maturation, genes involved in metabolism and cell cycle were consistent regulated among organs. These trends were expected as cell cycle rate decreases when organ size reaches homeostasis. In addition, the transition from glycolysis to fatty acid oxidation after birth has been well described in the heart (Murphy and Chen et al. 2021).

We took multiple approaches in gene regulatory network reconstruction to infer key upstream regulators. Briefly, we used fuzzy clustering to select genes that trended up or down in expression from early embryonic to adult timepoints to select genes that become more expressed or repressed during maturation. We see that the overlapping genes in the downregulated cluster were related to mRNA splicing, possibly indicating that a spliceosome construct associated with immature tissues is down regulated.

Other groups have studied comparisons between species such as human and mouse (Anzai et al. 2020), mouse and zebrafish (Irie et al. 2011), and human and macaque (Cardoso-Moreira et al. 2019). We decided to focus on the mouse as this built the most complete developmental time series. Since our meta-analysis used microarray datasets, further validation using RNA-seq datasets could support our findings. Single cell RNA-seq in particular could allow analysis with increased granularity and overcome confounders including the heterogeneity of maturation as shown earlier (Murphy et al. 2021).

Future studies are needed to biologically validate these findings. For example, increasing expression of the positively enriched upstream regulators in stem cell-derived cardiomyocytes, hepatocytes, or neurons may promote their functional maturation. We note that activating the expression of PGC1α/PPAR, a regulator of metabolism and one of the positively enriched upstream regulators identified by IPA, indeed enhances cardiomyocyte maturation (Murphy et al. 2021). It remains to be seen if activating such metabolic regulators can enhance the maturation of other stem cell-derived cell types. Their maturation status can also be quantified by use of transcriptomic analysis (Kannan et al 2020, Uosaki et al. 2015) to aid this evaluation. This may help alleviate the maturation issue in the stem cell field and hasten the translation to stem cell-based modeling and therapies (Murphy and Chen et al. 2021).

## Methods

### Dataset compilation

Affymetrix Mouse 430 2.0 microarray datasets of whole brain, liver, kidney, and hearts were compiled and curated from Gene Expression Omnibus (GEO) as done previously (Uosaki et al. 2015). The datasets covered all major developmental stages from embryonic to adult in all four organs. Datasets corresponding to wild-type and nontreated conditions were subsequently subsetted and clustered according to early embryonic, mid embryonic, late embryonic, neonatal, young adult, and adult timepoints. In total, 479 brain, 147 liver, 142 kidney, and 212 heart datasets that fit selection criteria were analyzed. Data was analyzed in R using custom scripts.

### Principal Component Analysis

Principal Component Analysis was run on the full probeset expression data using the prcomp() function. Probeset expression data was converted to gene expression data by selecting the probes with the greatest interquartile range of expression. Differentially expressed genes across each stage in each organ were identified using limma. Genes with a log fold change > 2 and a p value < .01 were analyzed using Ingenuity Pathway Analysis to infer upstream regulators.

### Fuzzy clustering

Fuzzy clustering of gene expression data was performed using Mfuzz (Kumar et al. 2007, Futschik et al. 2005). The number of clusters were determined by iteratively increasing the number of clusters until the correlation between any two clusters exceeded 0.85.

### Gene regulatory network

Weighted Gene Co-Expression Network Analysis was also performed on the gene expression data (Langfelder et al. 2008). Organ-specific correlation networks were constructed using data across all time points, using a signed network with biweight midcorrelation, a soft thresholding power of 12, and a maximum block size of 21,000. Using developmental stage as an external trait, the modules that were most positively and negatively correlated with development and maturation were identified based on the correlation and associated p-value. The top 3000 genes with highest module membership values and the top 300 genes with highest intramodular connectivity scores were selected for each module positively and negatively associated with maturation in each organ. A consensus network was also constructed using data from all four organs.

### Ingenuity Pathway Analysis

Ingenuity Pathway Analysis was performed by identifying differentially expressed genes between each stage of organ development. This differential gene expression information for each organ was used for network reconstruction via Ingenuity Pathway Analysis in order to identify upstream regulators activated and inactivated between each stage of development for each organ. Ggplot2 R package was used to generate (Wickham 2016).

## References

Agostini M, Romeo F, Inoue S, Niklison-Chirou MV, Elia AJ, Dinsdale D, Morone N, Knight RA, Mak TW, Melino G. Metabolic reprogramming during neuronal differentiation. Cell Death Differ. 2016 Sep 1;23(9):1502–14.

Anzai T, Yamagata T, Uosaki H. Comparative Transcriptome Landscape of Mouse and Human Hearts. Front Cell Dev Biol. 2020 Apr 22;8:268.

Cardoso-Moreira M, Halbert J, Valloton D, Velten B, Chen C, Shao Y, Liechti A, Ascenção K, Rummel C, Ovchinnikova S, Mazin PV, Xenarios I, Harshman K, Mort M, Cooper DN, Sandi C, Soares MJ, Ferreira PG, Afonso S, Carneiro M, Turner JMA, VandeBerg JL, Fallahshahroudi A, Jensen P, Behr R, Lisgo S, Lindsay S, Khaitovich P, Huber W, Baker J, Anders S, Zhang YE, Kaessmann H. Gene expression across mammalian organ development. Nature. 2019 Jul;571(7766):505–509.

Eisenberg E, Levanon EY. Human housekeeping genes, revisited. Trends Genet. 2013 Oct;29(10):569–74.

Futschik M, Carlisle B. “Noise robust clustering of gene expression time-course data.” Journal of Bioinformatics and Computational Biology. 2005 965–988.

Goss RJ. Similarities and differences between mechanisms of organ and tissue growth regulation. Proc Nutr Soc. 1990 Oct;49(3):437–42.

He Z, Yu Q. Identification and characterization of functional modules reflecting transcriptome transition during human neuron maturation. BMC Genomics. 2018 Apr 17;19(1):262.

Irie, Naoki, and Shigeru Kuratani. “Comparative transcriptome analysis reveals vertebrate phylotypic period during organogenesis.” Nature communications vol. 2 (2011): 248.

Kannan S, Farid M, Lin BL, Miyamoto M, Kwon C. Transcriptomic entropy benchmarks stem cell-derived cardiomyocyte maturation against endogenous tissue at single cell level. bioRxiv. 2020.

Kojima Y, Tam OH, Tam PP. Timing of developmental events in the early mouse embryo. Semin Cell Dev Biol. 2014 Oct;34:65–75.

Kumar, L., E Futschik, M. Mfuzz: a software package for soft clustering of microarray data. Bioinformation. 2007 2, 5–7.

Langfelder P, Horvath S. “WGCNA: an R package for weighted correlation network analysis.” BMC Bioinformatics. 2008 559.

Liang, S., Li, Y., Be, X., Howes, S., Liu, W. Detecting and profiling tissue-selective genes. Physiological genomics. 2006 26:158–162.

Lopaschuk GD, Jaswal JS. Energy metabolic phenotype of the cardiomyocyte during development, differentiation, and postnatal maturation. J Cardiovasc Pharmacol. 2010 56:130–40.

Mitiku N, Baker JC. Genomic analysis of gastrulation and organogenesis in the mouse. Dev Cell. 2007 Dec;13(6):897–907.

Murphy SA, Chen EZ, Tung L, Boheler KR, Kwon C. Maturing heart muscle cells: Mechanisms and transcriptomic insights. Semin Cell Dev Biol. 2021 May 2:S1084-9521(21)00092-6.

Murphy SA, Miyamoto M, Kervadec A, Kannan S, Tampakakis E, Kambhampati S, Lin BL, Paek S, Andersen P, Lee DI, Zhu R, An SS, Kass DA, Uosaki H, Colas AR, Kwon C. PGC1/PPAR drive cardiomyocyte maturation at single cell level via YAP1 and SF3B2. Nat Commun. 2021 Mar 12;12(1):1648.

Nishikawa T, Bellance N, Damm A, Bing H, Zhu Z, Handa K, Yovchev MI, Sehgal V, Moss TJ, Oertel M, Ram PT, Pipinos II, Soto-Gutierrez A, Fox IJ, Nagrath D. A switch in the source of ATP production and a loss in capacity to perform glycolysis are hallmarks of hepatocyte failure in advance liver disease. J Hepatol. 2014 Jun;60(6):1203–11.

Oh Y, Cho GS, Li Z, Hong I, Zhu R, Kim MJ, Kim YJ, Tampakakis E, Tung L, Huganir R, Dong X, Kwon C, Lee G. Functional Coupling with Cardiac Muscle Promotes Maturation of hPSC-Derived Sympathetic Neurons. Cell Stem Cell. 2016 Jul 7;19(1):95–106.

Qin Y, Pan J, Cai M, Yao L, Ji Z. Pattern Genes Suggest Functional Connectivity of Organs. Sci Rep. 2016 May 26;6:26501.

Robb L, Tam PP. Gastrula organiser and embryonic patterning in the mouse. Semin Cell Dev Biol. 2004 Oct;15(5):543–54.

Shyh-Chang N, Ng HH. The metabolic programming of stem cells. Genes Dev. 2017 Feb 15;31(4):336–346.

Tam PP, Behringer RR. Mouse gastrulation: the formation of a mammalian body plan. Mech Dev. 1997 Nov;68(1-2):3–25.

Takaoka K, Hamada H. Cell fate decisions and axis determination in the early mouse embryo. Development. 2012 Jan;139(1):3–14.

Tumaneng K, Russell RC, Guan KL. Organ Size Control by Hippo and TOR Pathways. Curr Biol. 2012. 22(9):R368–R379.

Uosaki H, Cahan P, Lee DI, et al. Transcriptional Landscape of Cardiomyocyte Maturation. Cell Rep. 2015 Nov;13(8):1705–1716.

Wang, ZY., Leushkin, E., Liechti, A. et al. Transcriptome and translatome co-evolution in mammals. Nature. 2020. 588, 642–647.

Warrington, J. A., Nair, A., Mahadevappa, M., Tsyganskaya, M. Comparison of human adult and fetal expression and identification of 535 housekeeping/maintenance genes. Physiol. Genomics. 2000. 2, 143–147.

Wickham H. Ggplot2: Elegant Graphics for Data Analysis. 2016. New York, NY: Springer-Verlag.

Xue L, Yi H, Huang Z, Shi YB, Li WX. Global gene expression during the human organogenesis: from transcription profiles to function predictions. Int J Biol Sci. 2011. 7(7):1068–76.

Yuan Q, Miao J, Yang Q, Fang L, Fang Y, Ding H, Zhou Y, Jiang L, Dai C, Zen K, Sun Q, Yang J. Role of pyruvate kinase M2-mediated metabolic reprogramming during podocyte differentiation. Cell Death Dis. 2020 May 11;11(5):355.

Zheng X, Boyer L, Jin M, Mertens J, Kim Y, Ma L, Ma L, Hamm M, Gage FH, Hunter T. Metabolic reprogramming during neuronal differentiation from aerobic glycolysis to neuronal oxidative phosphorylation. Elife. 2016 Jun 10;5:e13374.

